# Fibroblast-specific Deletion of *Yap/Taz* Impairs Mouse Postnatal Dermal Development by Suppressing Collagen Production and Deposition

**DOI:** 10.64898/2026.01.19.700422

**Authors:** Alexandre, Ava J Kim, Kirk C Hansen, Maxwell McCabe, Jun Young Kim, Zhaoping Qin, Zhaolin Zhang, Tianyuan He, Chunfang Guo, John J Voorhees, Gary J Fisher, Taihao Quan

## Abstract

The Hippo pathway effectors YAP and TAZ are key regulators of cell proliferation, apoptosis, and differentiation, thereby maintaining tissue homeostasis and controlling organ size. While their roles in epithelial tissues and cancer are well established, their role in dermal fibroblast extracellular matrix (ECM) regulation is less understood. Here, we investigated the role of *Yap/Taz* during postnatal skin dermis development. During postnatal growth, mouse skin steadily grows and undergoes significant surface expansion. Postnatal deletion of *Yap/Taz* in dermal fibroblasts, the primary cells responsible for dermal ECM homeostasis, significantly impaired dermal maturation, as evidenced by marked deficiencies in collagen synthesis and deposition. Isolated fibroblasts from *Yap/Taz* knockout mice showed reduced proliferation and diminished expression of *Yap/Taz* target genes (*Ccn2*, *Col1a1*), which were rescued by reintroduction of active *Yap/Taz*. RNA-seq, and spatial transcriptomics and proteomics of *Yap/Taz* knockout skin revealed substantial downregulation of matrisome genes, including type I (*Col1a1, Col1a2)* and type III (*Col1a3*) collagens, which together constitute more than 90% of the skin’s collagen content. These findings demonstrate that *Yap/Taz* are essential for dermal ECM homeostasis, highlighting their therapeutic potential in skin regeneration, fibrosis, and aging-related ECM decline.

## INTRODUCTION

The Hippo signaling pathway is a fundamental regulator of cell proliferation, apoptosis, and differentiation, ensuring tissue homeostasis and proper organ size (Moya and Halder 2019). Central to this pathway are its key downstream effectors, Yes-associated protein (YAP) and transcriptional coactivator with PDZ-binding motif (TAZ), which act as transcriptional regulators modulating gene expression in response to mechanical and biochemical signals.

Dysregulation of YAP/TAZ has been implicated in numerous human diseases, including cancer, fibrosis, and neurodegenerative disorders, highlighting their significance in pathophysiology (Dey et al. 2020). While extensive research has elucidated their roles in epithelial tissues and oncogenesis, the involvement of YAP/TAZ in regulation of the skin dermal extracellular matrix (ECM) remains largely unexplored. Given that the ECM provides structural and functional support to skin, understanding the impact of YAP/TAZ on ECM composition and remodeling is important for advancing knowledge of skin ECM biology and related disorders.

Although mouse skin, including hair follicles (HFs), forms during the prenatal period, it remains structurally immature at birth and undergoes extensive postnatal remodeling to establish mature skin architecture. This development is tightly regulated and involves interactions between epidermal, dermal, and immune components. Moreover, dermal maturation is influenced by the dynamic nature of skin, particularly fluctuations in thickness during the hair cycle. Fully developed dermal architecture is essential for mechanical strength, barrier integrity, immune defense, and wound healing. Despite the rapid postnatal growth in mice, the mechanisms underlying dermal maturation during this period are not well understood. This process is largely driven by dermal fibroblasts, which synthesize and deposit collagen, the primary structural protein responsible for skin dermal maturation.

We previously reported that knockdown of YAP/TAZ suppresses collagen production (Qin et al. 2022) and impairs TGF-β/Smad signaling in dermal fibroblasts (Qin et al. 2018), the primary regulators of collagenous ECM production (Border and Noble 1994; Quan et al. 2010; Deng et al. 2024). These findings led us to hypothesize that knockout of *Yap/Taz* in dermal fibroblasts may impair postnatal dermal development by inhibiting collagen production. To test this hypothesis, we investigated the role of *Yap/Taz* in postnatal dermal development by generating mice with a conditional knockout of *Yap/Taz* in dermal fibroblasts, the main ECM-producing cells in the skin. Fibroblast-specific deletion of *Yap/Taz* significantly impaired dermal maturation, primarily due to reduced production and deposition of collagen-rich ECM. These findings highlight the critical role of YAP/TAZ in maintaining dermal ECM integrity and fibroblast function, with potential implications for therapies targeting skin aging, fibrosis, and regenerative disorders.

## RESULTS

### Mouse skin dermis postnatal maturation

Newborn mice are born naked, blind, and without visible ears. Since mice are typically weaned around 21 days after birth, those younger than 21 days are generally considered to be in the early postnatal period. While the growth of the epidermis during this time is well studied (Dekoninck et al. 2020), the postnatal expansion of the dermis remains less well understood. We observed that skin surface area had expanded exponentially by postnatal day 20 (P20), with approximately 6-fold greater surface area in mice at P20 compared to postnatal day 0 (P0) (Figure 1a). Interestingly, the number of total dermal fibroblasts remained unchanged during the postnatal stages from P0 to P20 (Figure 1b). This skin area expansion, along with the lack of change in total dermal fibroblasts, indicates a decrease in the density of dermal fibroblasts during postnatal development (Figure 1c).

**Figure 1.**
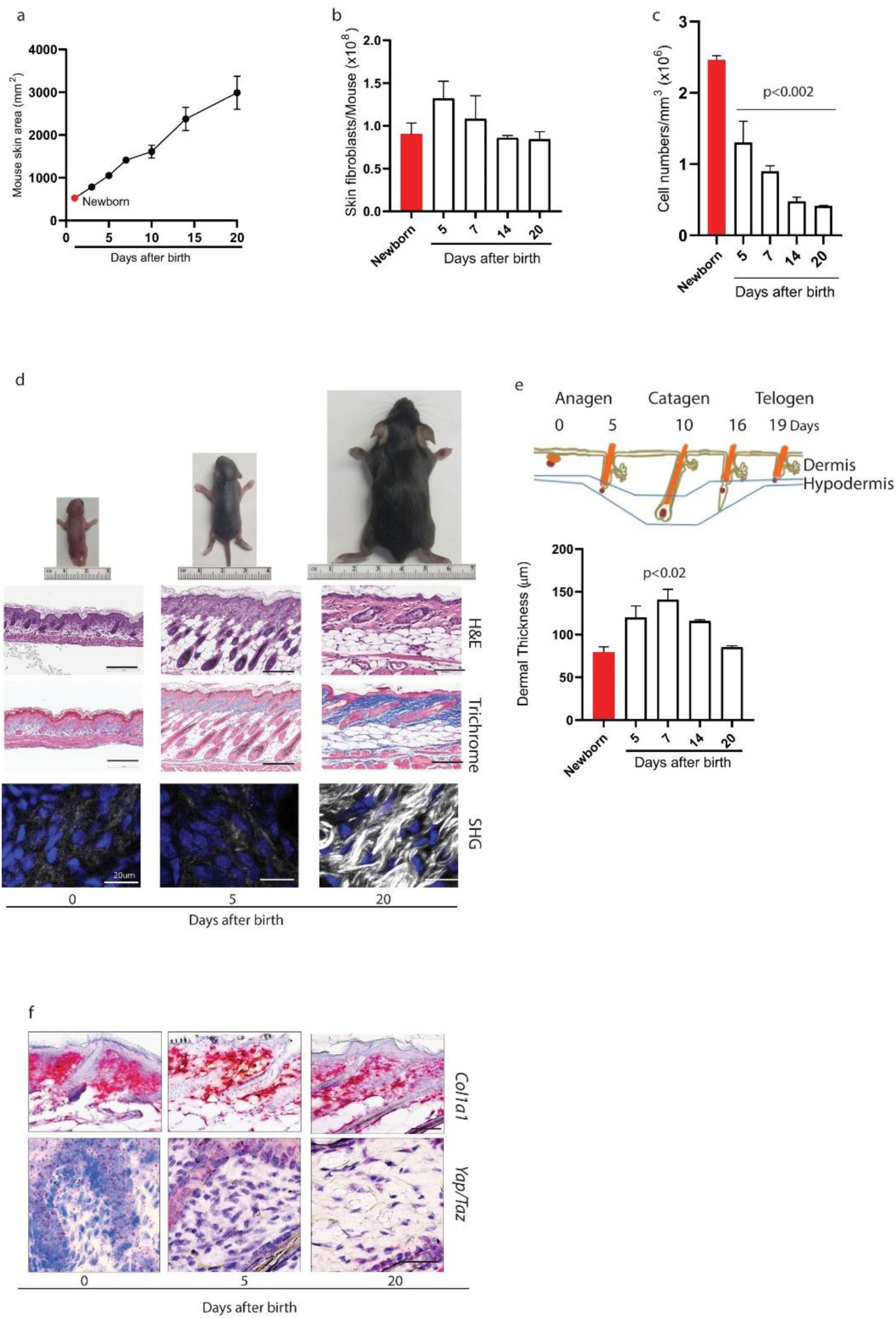
Postnatal maturation of the mouse skin dermis. Postnatal mouse skin area (**a**), dermal fibroblast number (**b**), and dermal fibroblast density (**c**) were quantified by ZEN Microscopy Software (ZEN Blue3, ZEISS). (**d**) Postnatal mouse skin histology: H&E (top) and trichrome middle), and SHG (bottom). (**e**) Postnatal mouse skin thickness was quantified by ZEN Microscopy Software (ZEN Blue3, ZEISS). (**f**) Postnatal mouse skin RAN scope® *in situ* hybridization: *Col-1a1* (top) and *Yap/Taz* (bottom). Data are presented as mean ± SEM. p-values are shown in the respective graphs. Scale bar = 200 µm (scale bar in SHG image = 20 µm).

Mouse skin development from P0 to P20 involves dramatic structural maturation (Figure 1d) as the skin transitions from a relatively immature status to a fully competent organ. At birth (P0), mouse skin is structurally immature, characterized by a notably thin dermis lacking mature collagen along with immature hair follicles and sebaceous glands (Figure 1d, left). By P5, the first synchronized hair cycle begins as hair follicles enter anagen (growth phase), marking an important developmental milestone. However, collagen deposition is not yet evident, suggesting that dermal collagen remains in an immature state at P5 (Figure 1d, middle). At P20, hair follicles begin to enter catagen (regression phase), completing the first synchronized hair cycle characteristic of early postnatal mouse development (Figure 1d, right). Most notably, the skin appears fully developed, with a mature epidermis, well-formed hair follicles, and dermal maturation. We also observed that the thickness of the dermis changed significantly with hair cycle progression (Figure 1d). The thickest dermis was observed in P7, in which HFs actively grow (anagen) (Figure 1d). Dermal thickness returns to its original (newborn/P0) level in P20 mice, in which HF regression is complete (telogen). Focusing on the dermis, we performed *Col1a1* in situ hybridization, as *Col1a1* is a major collagen component of the skin (Figure 1f). *Col1a1* transcription was highly expressed throughout the dermis from P0 to P20. In contrast, *Yap/Taz in situ* hybridization revealed a gradual decrease in expression with age, correlating with skin maturation. Together, postnatal skin development is marked by a significant expansion in surface area without an increase in total dermal fibroblast number, leading to a reduction in fibroblast density. Dermal thickness fluctuates with the hair cycle, peaking at P7 during anagen and returning to newborn levels by P20 in telogen. In newborn mice, the skin remains in an immature state, appearing fully developed by P20. *Col1a1* expression remains consistently high throughout dermal development, while *Yap/Taz* expression decreases with age, aligning with skin maturation.

### Role of *Yap/Taz* in mouse skin dermal postnatal maturation

To explore the role of *Yap/Taz* in mouse skin dermal maturation, we initially attempted double knockout *Yap* and *Taz* in dermal fibroblasts at embryonic stage by crossing *Yap/Taz* floxed (*Yap ^fl/fl^;Taz ^fl/fl^*) mice with stromal cell-specific Cre-driver mice (*Pdgfrα-Cre*) (Figure 2a). However, we observed that embryonic *Yap* and *Taz* knockout in fibroblasts (Figure 2b) resulted in embryonic lethality, primarily due placental underdevelopment (Figure 2c). Further investigation revealed that a fibroblast-specific *Yap* knockout alone is also embryonic lethal, while *Taz* knockout and other *Yap/Taz* heterozygous combinations are not (Table I).

**Figure 2.**
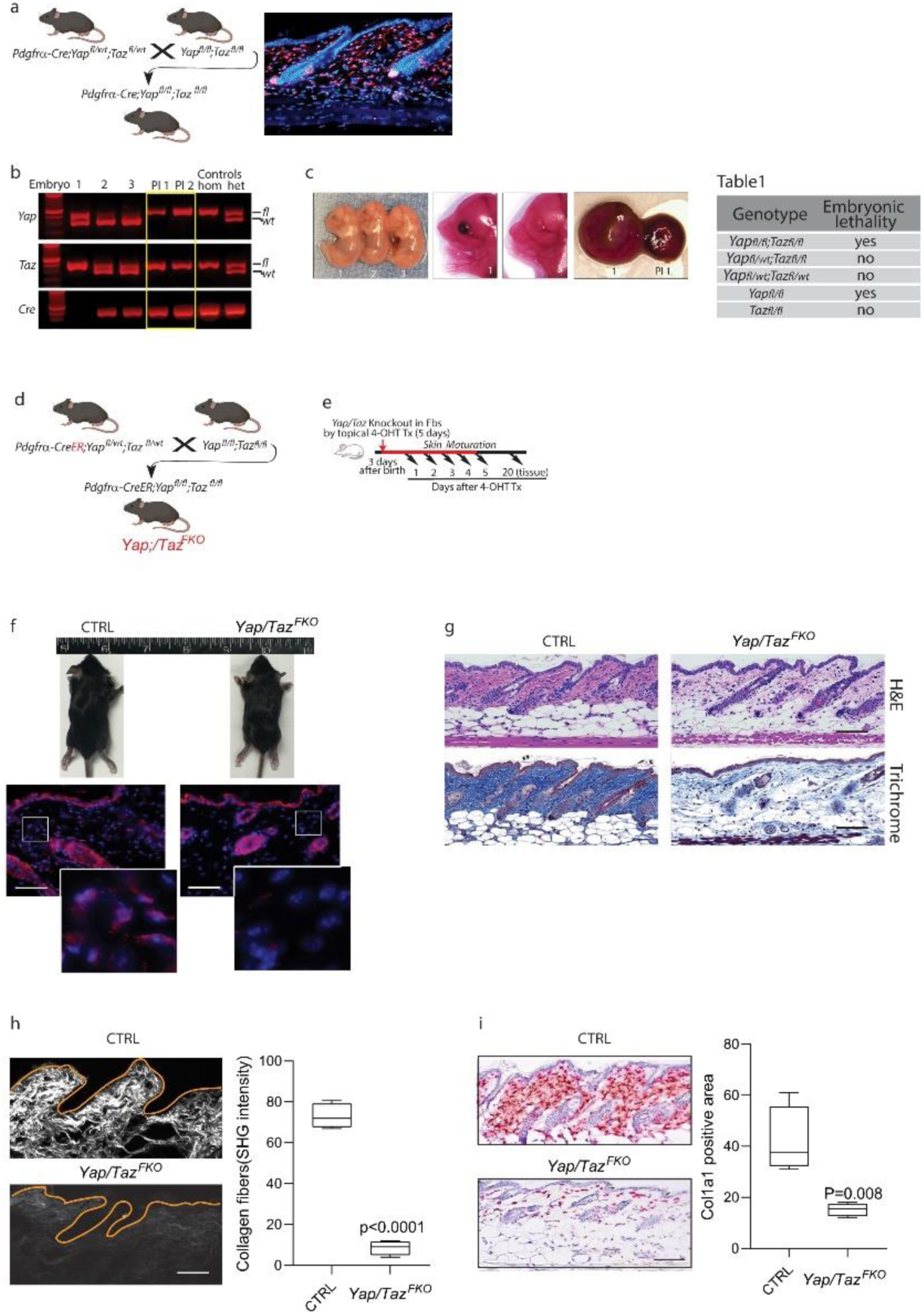
Role of *Yap/Taz* in mouse skin dermal maturation. (**a**) Schematic representation of embryonic fibroblast-specific *Yap* and *Taz* knockout mice (*pdgfra-Cre;Yap^f/f^;Taz^f/f^*). Image shown skin dermal fibroblast-specific staining of RFP driving by *Pdgfra* promoter in *ROSA26 Pdgfra-RFP* mice. (**b**) Embryo mouse genotype, with the yellow box highlighting the *Cre* positive *Yap/Taz* KO mice (*pdgfra-Cre;Yap^f/f^;Taz^f/f^*). (**c**) Image of embryo (E14.5) and placenta. Normal (left #1) and abnormal (Pl1, *pdgfra-Cre;Yap^f/f^;Taz^f/f^*) placentas were shown. (**d**) Schematic representation of postnatal inducible fibroblast-specific *Yap* and *Taz* KO mice (*Yap/Taz^FKO^*).(**e**) Schematic representation of topical 4-OH tamoxifen (10 mg/ml) treatment. (**f**) *Yap/Taz* immunohistology. No *Yap/Taz* staining is observed in dermal cells of *Yap/Taz* KO mice (*Yap/Taz^FKO^*), but staining is present in epidermal cells. (**g**) Fibroblast-specific *Yap/Taz* deletion impairs dermal collagen deposition. Top: H&E staining, bottom: trichrome staining. Note the reduced collagen fiber staining in both H&E and trichrome images. (**h**) Second harmonic generation (SHG) microscopy. Control mice (top) show thick, densely packed collagen bundles, which are absent in *Yap/Taz* KO mice (button, *Yap/Taz^FKO^*). (**i**) *Col1a1 in situ* hybridization (RNAscope™). Reduced *Col1a1* transcription is observed in *Yap/Taz* KO mice (*Yap/Taz^FKO^*). Quantification was performed using ImageJ software (version 1.5p). Data are presented as mean ± SEM. p-values are shown in the respective graphs. Scale bar=100 µm.

To overcome this, we generated mice with an inducible knockout of *Yap* and *Taz* in dermal fibroblasts after birth (*Pdgfra-CreER;Yap^fl/fl^;Taz^fl/fl^*, referred to herein as *Yap/Taz^FKO^*) (Figure 2d). To knockout *Yap/Taz* in dermal fibroblasts, P3 (postnatal day 3) mice and control sex-matched littermates were topically treated with 4-OH tamoxifen (10mg/ml) for five consecutive days, and back skin was harvested at P20 (Figure 2e). We confirmed dermal fibroblast-specific deletion *Yap/Taz* after topical 4-OH tamoxifen treatment (Figure 2f).

We found that fibroblast-specific deletion of *Yap/Taz* significantly impaired skin dermal maturation during the period prior to P20, evidenced by abolished production, deposition, and maturation of collagen fibrils (Figure 2g-i). Skin H&E and trichrome staining shows significant reduction of collagen bundles (Figure 2g, left). Second harmonic generation (SHG) microscopy, which provides high-resolution three-dimensional (3D) images of collagen fibers, indicated that collagen bundles in control mice appeared thick, densely packed, tightly interwoven, and evenly dispersed across the dermis (Figure 2h, top). In contrast, collagen bundles in *Yap/Taz^FKO^* mice were barely detectable (Figure 2h, bottom). Quantitative analysis showed a significant reduction in collagen bundle intensity in *Yap/Taz^FKO^* mice (Figure 2h, bar graph). Supporting the above data*, Col1a1* transcription is markedly reduced in *Yap/Taz^FKO^* mice (Figure 2i). These data indicate that *Yap/Taz* plays a critical role in skin dermal maturation by controlling dermal collagen synthesis and deposition.

### Impaired *Yap/Taz* function in *Yap/Taz^FKO^* mouse dermal fibroblasts

To investigate the functional consequences of *Yap/Taz* deletion, we isolated dermal fibroblasts from the skin of *Yap/Taz^FKO^* and control mice. First, we confirmed that *Yap/Taz* is highly expressed in control fibroblasts but markedly reduced in *Yap/Taz^FKO^* fibroblasts (Figure 3a). Next, we examined the expression of *Ccn2*, a well-established *Yap/Taz* target gene critically involved in collagen regulation (Quan et al. 2010; Quan et al. 2013; Qin et al. 2022). Our results reveal a marked decrease in *Ccn2* for *Yap/Taz^FKO^* fibroblasts at both the mRNA (Figure 3b) and protein levels (Figure 3c), indicating impaired *Yap/Taz* signaling. Notably, *Yap/Taz* knockout in dermal fibroblasts resulted in a substantial decrease in *Col1a1* mRNA expression (Figure 3d) and type I collagen protein levels (Figure 3e) in *Yap/Taz^FKO^* mice, indicating the pivotal role of *Yap/Taz* in regulating collagen production.

**Figure 3.**
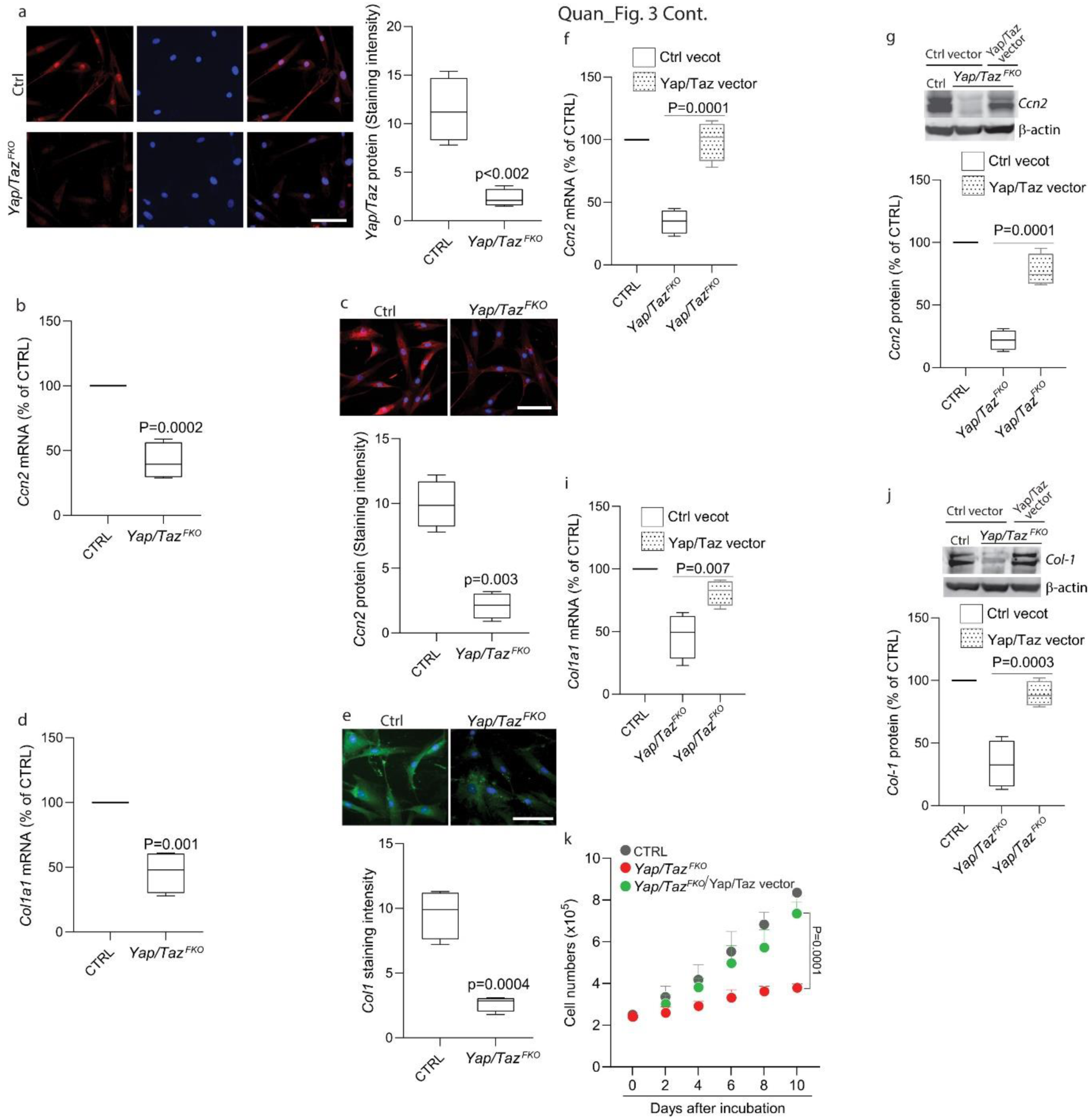
Impaired *Yap/Taz* function in *Yap/Taz^FKO^* mouse dermal fibroblasts. Dermal fibroblasts were isolated from the skin of *Yap/Taz* KO mice (*Yap/Taz^FKO^*) and control mice. (**a**) *Yap* and *Taz* immunostaining. Less *Yap/Taz* staining in the cells of *Yap/Taz* KO mice (*Yap/Taz^FKO^*), but staining is present in control (CTRL) cells. (**b**) *Ccn2* mRNA. (**c**) *Ccn2* protein immunostaining. (**d**) *Col1a1* mRNA. (**e**) Type I procollagen immunostaining. (**f**-**k**) Cells were transiently transfected with a YAP/TAZ mutant plasmid by electroporation. (**f**) *Ccn2* mRNA and (**g**) *Ccn2* protein Western blot. (**i**) *Col1a1* mRNA and (**j**) Type I collagen Western blot. (**k**) Cell proliferation. 2.5×10^5^cells were cultured in 60 mm plates for the indicated days and counted. mRNA and protein levels were determined by real-time PCR and Western blot analysis, respectively. mRNA levels were normalized to 36B4, a ribosomal protein used as an internal control. Protein levels were normalized to β-actin as loading control. Insects show representative Western blots. Results are expressed as mean ± SEM, N=4. p-values are shown in the respective graphs. Scale bar=100 µm.

Next, we asked whether restoring *Yap/Taz* expression in *Yap/Taz^FKO^*mouse dermal fibroblasts could rescue collagen expression. To test this, *Yap/Taz* expression was restored by introducing *Yap/Taz* mutant plasmids (see Materials & Methods for details), which promote constitutive *Yap/Taz* nuclear translocation (Chan et al., 2011, (Qin et al. 2022). Restoration of *Yap/Taz* expression almost completely rescued *Ccn2* mRNA (Figure 3f) and protein levels (Figure 3g), as well as *Col1a1* mRNA (Figure 3i) and type I collagen protein levels (Figure 3j).

Finally, we explored the impact of *Yap/Taz* knockout on dermal fibroblast proliferation. We found that *Yap/Taz* knockout significantly reduced dermal fibroblast proliferation (Figure 3k, red line). Notably, restoring *Yap/Taz* expression rescued cell proliferation (Figure 3k, green line). Together, these data suggest that *Yap/Taz* plays a critical role in dermal fibroblast collagen regulation and proliferation.

### Fibroblast-specific *Yap/Taz* knockout mouse transcriptome

Bulk RNA-seq analysis revealed that 2668 genes were differentially expressed (1466 upregulated, 1202 downregulated) in *Yap/Taz^FKO^*mice compared to non-transgenic control mice (Figure 4a, 4b). As expected, many collagens, including the major collagens in skin (*Col1a1, Col1a2, and Col1a3*), are significantly reduced (Figure 4c), whereas multiple matrix metalloproteases (MMPs) were upregulated in *Yap/Taz^FKO^* mice (Figure 4d).

**Figure 4.**
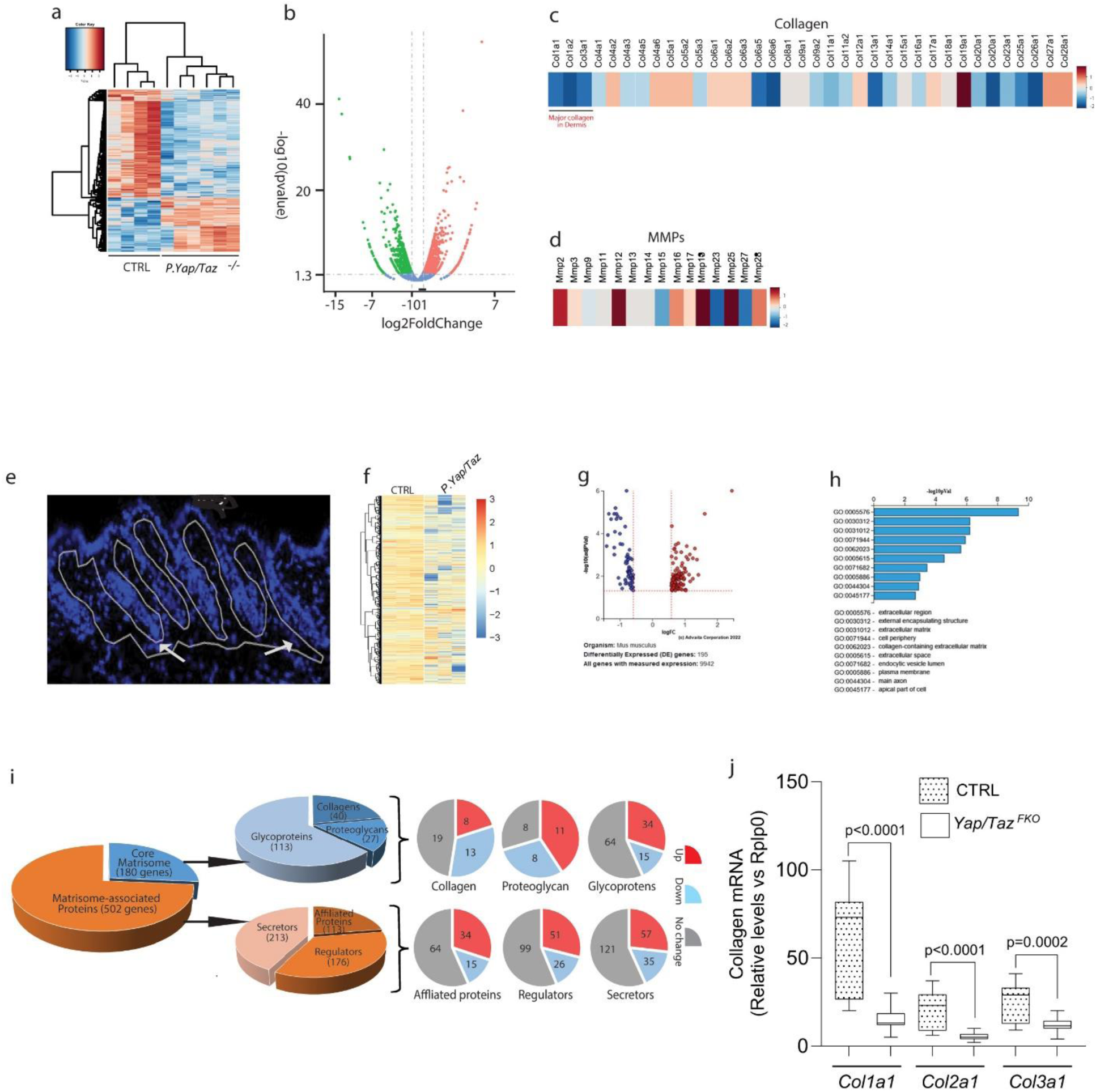
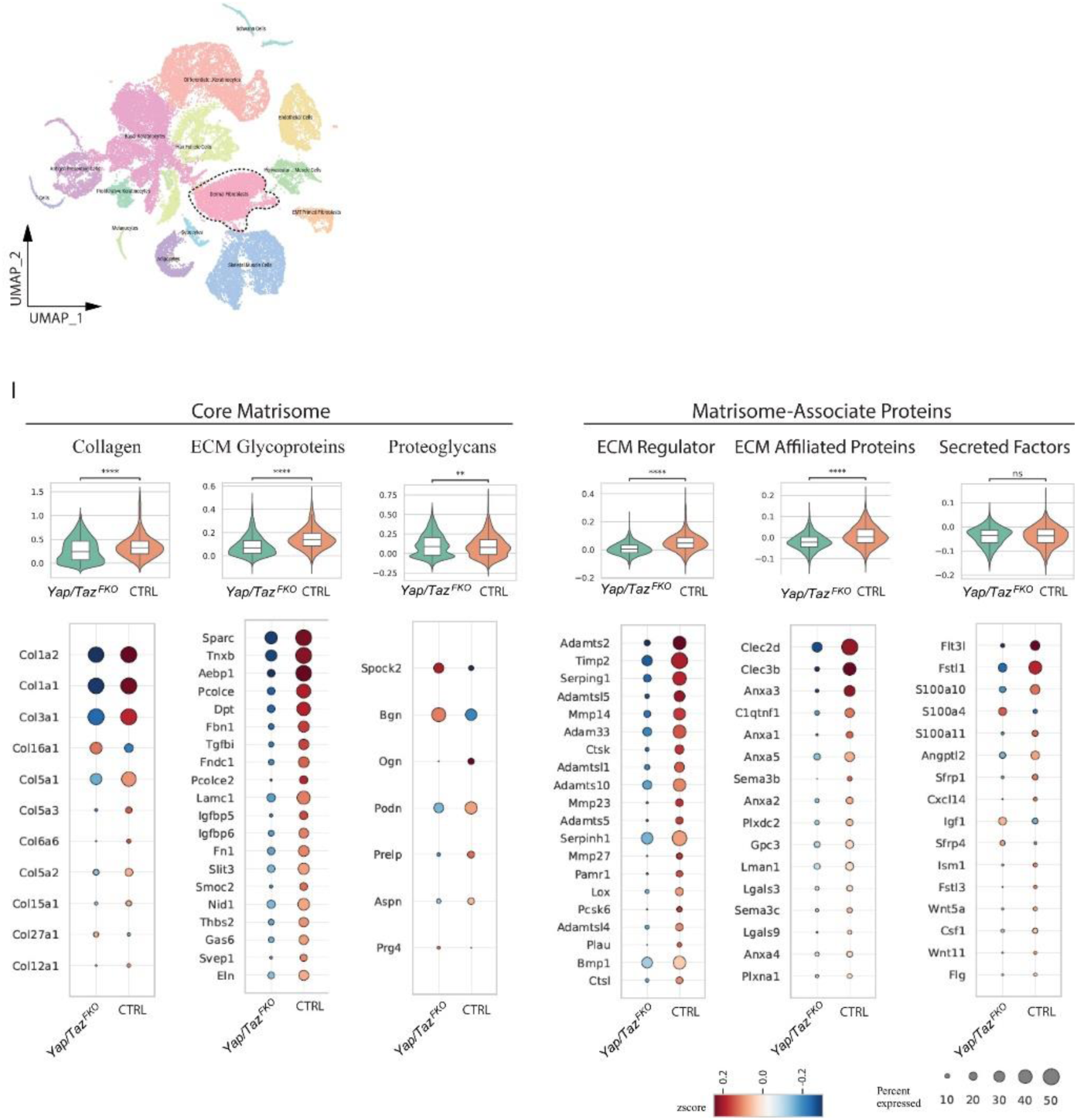
Fibroblast-specific *Yap/Taz* knockout mouse transcriptome. **(a-d)** Bulk RNA-seq. (**a**) Heatmap and (**b**) volcano plots showing the change of genes; total 2666 genes were differentially expressed (1466 upregulated, 1202 downregulated) in *Yap/Taz* KO mice (*Yap/Taz^FKO^*). (**c** and **d**) Heatmap showing significant downregulation of type I and other collagens (**c**) and upregulation of multiple *Mmps* in *Yap/Taz KO mice* (*Yap/Taz^FKO^*). (**e**-**J**) Spatial transcriptomics. Spatial transcriptomics was performed using the NanoString GeoMx® Digital Spatial Profiler. (**e**) Regions of interest (ROIs) within the interstitial dermis, where dermal fibroblasts reside (DAPI-stained images). (**f**) Heatmap. (**g**) volcano plot displays the distribution of the 195 most significantly differentially expressed genes. (**h**) GO terms shows genes involved in ECM function mostly altered in *Yap/Taz* KO mice (*Yap/Taz^FKO^*). (**i**) Matrisome profile. Total 682 matrisome genes were identified (180 core matrisome and 502 matrisome-associated genes). Note *Yap/Taz* knockout in dermal fibroblasts significantly altered the core matrisome signature (upper doughnut charts) and matrisome-associated genes (lower doughnut charts). (**j**) Type I (*Col1a1* and *Col1a2*) and type III (*Col1a3*) mRNA expression. Note *Col1a1, Col1a2,* and *Col1a3* were among the top three most downregulated DGEs. (**k**-**l**) scRNA-seq analysis reveals changes in the matrisome profile of aged dermal fibroblasts. (**k**) Uniform Manifold Approximation and Projection (UMAP) plot displaying eleven distinct cell clusters representing major skin cell types. (**l**) Violin plots illustrating transcriptomic alterations in core matrisome and matrisome-associated proteins and dot plots depicting the top DEGs. Results are expressed as mean ± SEM, N=3. *p-*values are shown in the respective graphs.

Given our focus on the ECM, we performed special sequencing (Figure 4e) to characterize the dermal matrisome, which comprises a diverse array of ECM components totaling 1,105 proteins (Hynes and Naba 2012; Naba et al. 2016). The matrisome is classified into two main groups: the core matrisome and matrisome-associated proteins. The core matrisome includes collagens, ECM glycoproteins, and proteoglycans, while matrisome-associated proteins encompass ECM-affiliated proteins, ECM regulators, and secreted factors. A heatmap illustrates differential gene expression in the dermal region of interest (ROI) between *Yap/Taz* knockout and control littermate mice (Figure 4f). A volcano plot illustrates the distribution of the 195 most significantly differentially expressed genes (Figure 4g). Notably, pathway analysis revealed significant representation of genes involved in ECM function (Figure 4h). We identified 682 matrisome genes, consisting of 180 core matrisome and 502 matrisome-associated genes (Figure 4I). Among the 180 core matrisome genes, we identified 40 collagens, 27 proteoglycans, and 113 glycoproteins. Additionally, *Yap/Taz* knockout in dermal fibroblasts significantly altered the proteome signature across both core matrisome and matrisome-associated protein groups (Figure 4i). Quantification of type I (*Col1a1, Col1a2*) and type III (*Col3a1*) collagen, the most abundant matrisome components in the skin, revealed a marked downregulation in *Yap/Taz* knockout mice (Figure 4j).

To further confirm our findings from bulk analysis of the matrisome transcriptome, we performed scRNA-seq. Dimensionality reduction using Uniform Manifold Approximation and Projection (UMAP) revealed 11 distinct clusters corresponding to major skin cell types, including keratinocytes, fibroblasts, skeletal muscle cells, adipocytes, hair follicle cells, endothelial cells, perivascular muscle cells, T cells, schwann cells, antigen-presenting cells, and sebocytes (Figure 4k). To further characterize the dermal matrisome signature, we again focused our analysis on dermal fibroblasts. The transcriptomic profiles of both core matrisome and matrisome-associated proteins were markedly downregulated in *Yap/Taz* knockout mice (Figure 4i). We highlighted the top DEGs in the dot plot, where most exhibited significantly in *Yap/Taz* knockout mice. Notably, *Col1a1* and *Col1a2*, encoding type I collagen, were among the top two downregulated DEGs, with *Col1a3* (type III collagen), the second most abundant collagen in skin, ranking within the top three downregulated genes (Figure 4i left dot plot). Collectively, these findings indicate that *Yap/Taz* knockout in dermal fibroblast profoundly alters the dermal matrisome transcriptional landscape, particularly through the significant downregulation of core matrisome collagens.

### ECM-focused proteomics in fibroblast-specific *Yap/Taz* knockout mouse skin

To complement the transcriptomic analysis and directly assess ECM composition, we performed ECM-focused proteomic profiling of mouse skin using a previously-established extraction workflow that separates soluble (cytosolic and loosely-bound ECM) and insoluble (structural ECM) protein fractions (McCabe et al. 2021). Skin samples from control and *Yap/Taz^FKO^* mice were fractionated, digested, and analyzed by LC-MS/MS. A total of 4,743 proteins were quantified, including 4,648 proteins in the soluble fraction and 1,058 in the insoluble ECM (iECM) fraction.

We identified 243 ECM-associated proteins by MatrisomeDB annotation, consisting of 105 core matrisome and 138 matrisome-associated proteins (Figure 5a). Among the 105 core matrisome proteins, we identified 22 collagens, 13 proteoglycans, and 70 ECM glycoproteins. Notably, fibroblast-specific *Yap/Taz* knockout significantly altered the abundance of both core matrisome and matrisome-associated proteins (Figure 5A), further supporting our transcriptomic results. Consistent with the known composition of dermal ECM, type I collagen (COL1A1/COL1A2) and type III collagen (COL3A1) were the most abundant proteins in the iECM fraction. Among the top 1000 most abundant proteins in the skin, collagen I accounted for ∼32% and collagen III for ∼9% of total MS signal intensity, highlighting their dominant structural role in the dermis. A decrease in type I collagen (∼20%) with *Yap/Taz* knockout was offset in the matrix by a significant increase in collagens COL4A3 (3.1 FC), COL6A1 (3.1 FC), COL6A2 (2.8 FC) and COL12A1 (1.8 FC). Several ECM glycoproteins were significantly decreased in *Yap/Taz* knockout mice, including SPARC (−3.4 FC), PCOLCE (−2.7), MFAP5 −1.7 FC), and several increased, including osteopontin (SPP1, 10.1 FC), basement membrane component LAMC3 (17 FC), FN1 (2.6 FC), fibrinogen (FGA,FGB,FGG 5.75 FC), VTN (4.0 FC), TNC (3.2 FC), POSTN (3.2 FC). Proteoglycans that decreased included OGN (−2.0 FC), while we conversely observed increases in BGN (2.2 FC), VCAN (2.3) and PRG4 (2.3).

**Figure 5.**
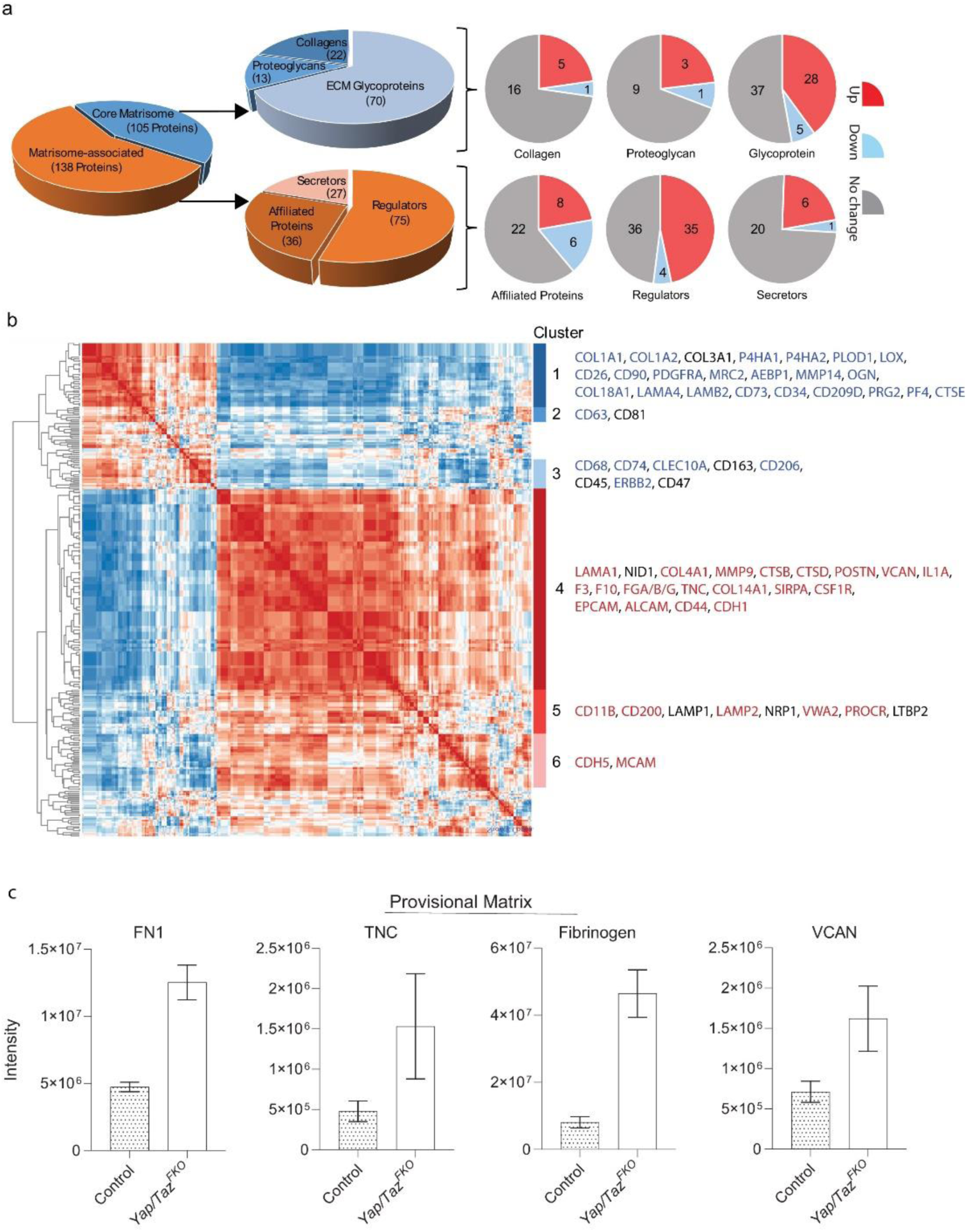
ECM-focused proteomics in in fibroblast-specific *Yap/Taz* knockout mouse skin. ECM-focused proteomic analysis reveals shift in cell and matrix signatures in fibroblast-specific *Yap/Taz* knockout mouse skin. **a**) Pie charts display identified matrisome proteins by category (left) and subcategory (center left). Pie charts on right display proteins significantly up (red; p < 0.05, FC > 1) and downregulated (blue; p < 0.05, FC < 1) in *Yap/Taz* KO samples by matrisome subcategory. Fold changes were calculated as average intensity for *Yap/Taz* KO samples divided by average intensity for control samples. **b**) Heatmap displaying Spearman correlation between all identified matrisome and cell marker proteins. Proteins contained in each labeled cluster are listed on the right, with proteins upregulated (red; FC > 1.15) and downregulated (blue; FC < 0.85) in *Yap/Taz* KO samples indicated by color. **c**) Bar plots displaying detected intensity for provisional matrix proteins significantly elevated in Yap/Taz KO samples. Error bars represent standard deviation.

Unsupervised clustering of differentially abundant stromal proteins across control and *Yap/Taz^FKO^* samples revealed distinct expression patterns, mirroring transcriptomic trends (Figure 5b). Cluster 1, enriched in control samples, included canonical fibroblast ECM components such as COL1A1, COL1A2, P4HA2 (6.6 FC), P4HA1 (1.9 FC), PLOD1(2.2 FC), LOX (1.9 FC), and SERPINH1 (4.1 FC), indicating preserved collagen biosynthesis and matrix cross-linking. This cluster also displayed a strong fibroblast/mesenchymal component, additionally containing DPP4 (CD26), PDGFRA, THY1 (CD90), MRC2, AEBP, MMP14, and OGN. A subset of endothelial-associated proteins was also present, including COL18A1, LAMA4, LAMB2, and NT5E (CD73), along with CD34, which marks dermal fibroblast subsets and hematopoietic progenitors. Immune associated proteins in this cluster included CD209D (dendritic), PRG2 (eosinophils), PF4 (platelets), and CTSE (macrophages). Clusters 2 and 3, partially elevated in control mice, included proteins associated with M2-like macrophage presence (CD68, CD74, CLEC10A, CD206). These clusters also featured an immunoregulatory and wound resolution signature, including proteins such as CD45, ERBB2, and CD47. Cluster 4 includes proteins that increase in the *Yap/Taz^FKO^*proteome and is characterized by elevated levels of basement membrane proteins (LAMA1, COL4A1), inflammatory ECM modifiers (MMP9, CTSB, CTSD, PRSS), fibroblast-to-inflammatory transition markers (POSTN, S100A8/A9, VCAN, IL1A, IL1RN), acute phase and coagulation proteins (F3, F10, FGA/B/G), and provisional matrix proteins (VCAN, TNC, COL14A1), consistent with a shift toward an injury- or regeneration-associated ECM. Provisional matrix proteins FN1, TNC, fibrinogen, and VCAN are all significantly elevated in *Yap/Taz^FKO^*samples (Figure 5c), further supporting this compositional shift. Cell surface markers in this cluster include a macrophage component (SIRPA, CSF1R, CD93) and a strong epithelial or endothelial component (EPCAM, ALCAM, PECAM1, CD44, CDH1). Clusters 5 and 6 are enriched in immune response proteins (SIGLEC1, ITGAM (CD11B), Angiogenesis-associated proteins (NRP1, VWA2, CDH5, PROCR, MCAM) and latent TGFβ regulators (SERPINB2, SERPIN1E).

A volcano plot of protein abundance differences between control and *Yap/Taz^FKO^* samples is provided in the Supplemental Data (Supplemental Figure 1). Together, these proteomic results reinforce the conclusion that YAP/TAZ are essential regulators of dermal ECM composition and fibroblast matrix production. Loss of YAP/TAZ not only diminishes collagen output but also drives a qualitative shift in matrix architecture and cell signatures at the protein level.

## DISCUSSION

The laboratory mouse serves as a fundamental animal model for investigating skin biology and disease. While embryonic skin development has been well characterized (Hu et al. 2018), the maturation of the skin during postnatal stages remains relatively understudied. In newborn mice, the dermis is immature and undergoes significant changes during early postnatal development (∼P20). We demonstrate in this study that, during the early postnatal period, the dermis undergoes rapid expansion without a corresponding increase in total fibroblast number, resulting in decreased fibroblast density. We also show that dermal thickness varies significantly with the hair cycle: it is greatest during the anagen phase at P7 and returns to a newborn-like thickness during the telogen phase at P20. At birth (P0), mouse skin is immature, with a thin dermis, undeveloped collagen, and immature hair follicles and sebaceous glands. By P5, hair follicles enter the growth phase (anagen), starting the first synchronized hair cycle, but dermal collagen is still immature. By P20, as hair follicles begin regression (catagen), the first hair cycle ends, and the skin appears fully developed with a mature epidermis, mature hair follicles, and well-formed dermal collagen. Collagen plays a crucial role beyond mere structural support in early postnatal skin development, actively influencing tissue architecture, cell behavior, and developmental signaling. These functions are vital for the formation of fully functional and resilient skin.

Dermal fibroblasts are responsible for the production of the collagenous ECM. To investigate the role of *Yap/Taz* in postnatal dermal maturation, we selectively knocked out these genes in dermal fibroblasts. Initial attempts to delete *Yap/Taz* at the embryonic stage resulted in lethality due to placental defects, highlighting the essential function of *Yap/Taz* in early development. To circumvent this, we created an inducible postnatal *Yap/Taz* knockout model (*Yap/Taz^FKO^*). Key findings indicate that *Yap/Taz* deletion in fibroblasts severely disrupts dermal maturation by markedly inhibiting collagen production and deposition. SHG microscopy showed that normal collagen networks, which are thick and densely packed, were largely absent in *Yap/Taz* knockout mice. *Col1a1* transcription, a critical marker for collagen production, was significantly downregulated in fibroblast-specific *Yap/Taz*-deficient mice. These results demonstrate that *Yap/Taz* are crucial regulators of dermal fibroblast function, particularly in collagen synthesis and deposition, which are essential for skin dermal maturation.

By isolating fibroblasts from *Yap/Taz* knockout mice, we provide compelling evidence that *Yap/Taz* deletion leads to severe functional impairments in fibroblasts, particularly in collagen regulation and cell proliferation. Loss of *Yap/Taz* in dermal fibroblasts results in a marked reduction in *Ccn2,* a key *Yap/Taz* target gene involved in collagen regulation (Qin et al. 2022). This downregulation correlates with a significant decrease in *Col1a1* expression and type I collagen production, confirming that *Yap/Taz* is essential for maintaining fibroblast-driven collagen synthesis. Interestingly, reintroducing *Yap/Taz* expression in knockout fibroblasts restores normal levels of *Ccn2, Col1a1*, and type I collagen. This not only validates the direct role of *Yap/Taz* in collagen regulation but also suggests potential therapeutic strategies for skin disorders where collagen synthesis is impaired. In addition to collagen synthesis, our study highlights the importance of *Yap/Taz* in fibroblast proliferation, with knockout cells showing significantly reduced proliferation rates. The rescue experiment further supports this, as *Yap/Taz* re-expression reinstates normal proliferation levels. This finding suggests that *Yap/Taz* contributes to dermal homeostasis by maintaining an adequate fibroblast population, which is essential for skin repair and maintenance. These findings establish *Yap/Taz* as central regulators of dermal fibroblast function, influencing both collagen production and cell proliferation. The ability to restore collagen synthesis and cell growth by reintroducing *Yap/Taz* further highlights their therapeutic potential in skin regeneration and wound healing. For instance, we previously reported that enhancing dermal fibroblast spreading and mechanical support in aged (80+ years old) human skin *in vivo*, through injection of cross-linked hyaluronic acid (a dermal filler), induces a more “youthful” fibroblast phenotype by improved cell spreading and increased mechanical tension (Quan et al. 2013). YAP/TAZ activity is known to be regulated by cell shape and mechanical cues independently of the Hippo pathway (Dupont et al. 2011; Panciera et al. 2017). These findings suggest that promoting fibroblast spreading and mechanical tension in aged skin may restore diminished YAP/TAZ expression in the dermis (Xiang et al. 2021; Qin et al. 2022; Kim and Quan 2024). Thus, restoring YAP/TAZ function represents a promising therapeutic strategy for reversing dermal aging. Future studies should explore the downstream signaling pathways regulated by YAP/TAZ and assess their therapeutic potential in treating ECM-related skin disorders such as fibrosis, impaired wound healing, and age-related dermal atrophy.

It is intriguing how YAP/TAZ regulate collagen gene expression as it is unlikely that YAP/TAZ directly regulate collagen gene transcription. One plausible mechanism is through their downstream target CCN2, a well-established and bona fide YAP/TAZ target, which plays a critical role in collagen regulation (Qin et al. 2022). Another possibility involves the TGF-β/Smad signaling pathway, a major pathway regulating collagen synthesis (Quan et al. 2010; Deng et al. 2024). We previously reported that knockdown of YAP/TAZ in human skin dermal fibroblasts impairs TGF-β/Smad signaling by induction of Smad7 (Qin et al. 2018). Taking together with our current findings, these data suggest that YAP/TAZ function as essential regulators of collagen production, acting through CCN2 and in coordination with TGF-β/Smad signaling.

Our special and single-cell transcriptomic analyses provide compelling evidence that fibroblast-specific *Yap/Taz* deletion profoundly reprograms the dermal matrisome gene expression. The broad downregulation of key structural components, most notably, type I and type III collagens (*Col1a1, Col1a2, Col3a1*), was a defining feature of the *Yap/Taz* knockout phenotype, as highlighted by both bulk and special RNA-seq as well as scRNA-seq differential gene expression analyses. This reduction in collagen gene expression was accompanied by the upregulation of several MMPs, suggesting increased ECM remodeling activity. Matrisome-focused analysis further revealed that both the core matrisome and matrisome-associated genes were altered, with a marked bias towards downregulation. Dimensionality reduction and cluster analysis confirmed that fibroblasts, the principal matrisome regulators, were chiefly responsible for these transcriptional changes. Collectively, these findings highlight YAP/TAZ as pivotal transcriptional regulators of dermal ECM homeostasis, particularly through maintenance of the collagen-rich core matrisome signature. Disruption of YAP/TAZ function in fibroblasts thus leads to significant impairment of ECM gene networks, with potential implications for understanding skin integrity, wound healing, and fibrotic disease mechanisms.

Fibroblast-specific loss of Yap/Taz not only suppresses collagen gene expression but also drives a profound reprogramming of dermal ECM composition and organization, resulting in a provisional, injury-associated matrix state that closely parallels and functionally supports the identified transcriptomic matrisome signature. ECM-focused proteomics confirmed that *Yap/Taz* knockout leads to a broad reduction of canonical fibroblast-derived structural ECM components, including decreased type I collagen and key collagen-associated proteins such as SPARC, PCOLCE, MFAP5, and OGN. These changes align with the strong downregulation of core matrisome collagens and collagen-modifying enzymes observed in bulk, spatial, and single-cell transcriptomic datasets.

Importantly, despite the continued presence of fibrillar collagens at the protein level, second harmonic generation (SHG) microscopy demonstrated an almost complete loss of organized collagen signal in Yap/Taz-deficient skin, indicating a dramatic disruption of higher-order collagen architecture. In control dermis, SHG revealed thick, densely packed, and highly ordered collagen bundles characteristic of a mature, load bearing fibrillar matrix. In contrast, Yap/Taz knockout skin exhibited minimal SHG signal, consistent with a failure to assemble, align, or stabilize collagen fibrils into an organized supramolecular network. Together, these findings indicate that Yap/Taz loss compromises not only collagen abundance but also the structural organization and physical integrity of the dermal ECM, uncoupling collagen presence from collagen function.

Unsupervised proteomic clustering further identified a YAP/TAZ-dependent fibroblast ECM module encompassing COL1A1, COL1A2, P4HA1/2, PLOD1, LOX, SERPINH1, DPP4, PDGFRA, THY1, and MMP14, that is selectively enriched in control skin. This module provides protein-level evidence that YAP/TAZ activity sustains a collagen-rich, cross-linked, and mechanically coherent dermal matrix, as well as a mesenchymal fibroblast identity congruent with the transcriptional matrisome profile. In contrast, Yap/Taz-deficient skin exhibited coordinated increases in basement membrane and provisional matrix proteins, including COL4A3, COL6A1/2, COL12A1, FN1, TNC, POSTN, fibrinogen chains (FGA, FGB, FGG), VCAN, BGN, PRG4, SPP1, and LAMC3, reflecting a qualitative shift away from a collagen I-dominant fibrillar ECM toward an injury- or regeneration-associated matrix state.

Taken together, the integration of ECM proteomics with SHG imaging and transcriptomic profiling demonstrates that YAP/TAZ regulate both the molecular composition and the supramolecular architecture of the dermal ECM. While some fibrillar collagen persists in the absence of Yap/Taz, its failure to assemble into an organized, SHG-detectable network indicates a loss of functional matrix integrity. These data support a model in which YAP/TAZ are required not merely for collagen production, but for the coordinated biosynthesis, processing, and architectural maturation of the collagenous ECM that underpins postnatal dermal development. Loss of Yap/Taz thus traps the dermis in a remodeling-prone, mechanically deficient ECM state that likely contributes to impaired tissue maturation, altered fibroblast behavior, and dysregulated repair responses.

In summary, fibroblast-specific deletion of *Yap/Taz* in mouse skin impairs dermal maturation by significantly downregulating matrisome gene expression. These findings indicate that YAP/TAZ are critical for maintaining dermal ECM homeostasis, underscoring their therapeutic potential in skin regeneration, fibrosis, and age-related ECM decline.

## MATERIALS AND METHODS

### Materials

Dulbecco’s Modified Eagle’s Media (DMEM), fetal bovine serum, trypsin solution, penicillin, and streptomycin were purchased from Invitrogen Life Technology. Antibodies for Yap/Taz was purchased from Cell Signaling Technology, CCN2 was purchased from Santa Cruz Biotechnology. Col-1 antibody for Western blot was purchased from SouthernBiotech, and for immunohistology was purchased from Santa Cruz Biotechnology. Unless otherwise stated, all other reagents were purchased from Sigma Chemical Company.

### Generation of Pdgfra-CreER;Yap^f/lf^;Taz^fl/fl^ (Yap/Taz^FKO^) mice

The double *Yap flox/Taz flox* mutant mice were purchased from The Jackson Laboratory (Stock#: 030532). These mice possess *loxP* sites flanking exon 2 of both the *Yap1* and *Wwtr1* genes. Since *Yap flox/Taz flox* mice are mixed background (ICR, C57BL/6 and 129S), *Yap flox/Taz flox* mice were crossed with six or more generations of pure C57BL/6 mice (Stock #; 000664) to reduce the variability of the genetic background. *Yap flox/Taz flox* mutant mice were mated *Pdgfra-creER^TM^* transgenic mice (The Jackson Laboratory, Stock#:018280, C57BL/6 genetic background), with Cre expression under the control of a stromal cell-specific regulatory sequence from the *Pdgfra* (platelet derived growth factor receptor alpha) promoter (Figure 2d)(Horikawa et al. 2015). After several rounds of backcrossing, *Pdgfra-CreER;Yap^f/lf^;Taz^fl/fl^* (*Yap/Taz^FKO^*) mice were generated as the experimental cohort. To delete *Yap* and *Taz* in mouse skin dermal fibroblasts, *Yap/Taz^FKO^* mice and control sex-matched littermates were topically treated with 4-OH tamoxifen (10mg/ml) for five consecutive days, and back skin were harvested at P20 (Figure 2e). Tissue samples were fixed overnight at room temperature in neutral-buffered formalin and subsequently embedded in paraffin (Histoserv, Inc.). Additional samples were fixed in 4% paraformaldehyde (in PBS) at 4°C and embedded in OCT compound. All experiments performed with the mice followed the standards of care approved by the University of Michigan Unit for Laboratory Animal Medicine (ULAM). Protocols for mouse experimentation were approved by the University of Michigan Institutional Animal Care and Use Committee.

### Measurement of mouse back skin area, thickness, and fibroblasts density

Mice were euthanized with CO₂, and hair shafts on the dorsal skin were removed using electric clippers. To estimate trunk surface area, the length from the base of the ear to the base of the tail and the diameter at the level of the costal arch were measured. Trunk surface area (A) was calculated using the formula: 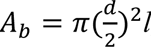*, where A_b_* – area, *d* – diameter, *l* – length. The Skin samples were analyzed using a Zeiss Axio Observer microscope with ZEN software. From each sample, 3-4 images at appropriate magnification (5× or 10×) were captured from three anatomical regions: cranial (near head), mid-trunk, and caudal (near tail). Dermal thickness was measured at 5-6 locations per image using a grid overlay and the length measurement tool in ZEN.

For fibroblast quantification, 3-4 regions of interfollicular dermis were selected per image. Fibroblasts within each region were counted using the counting tool, and the area of each counted region was recorded. All measurements were exported to Microsoft Excel for analysis. The density of fibroblasts per field (*D_f_*_2_) was calculated using the number of fibroblasts (*f*) in each area (*a*) normalized to 1mm^2^ 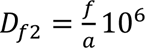, two-dimensional observation of fibroblasts density (*D_f_*_2_) was converted in three-dimensional observation 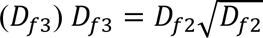. To estimate the total number of fibroblasts in the mouse trunk dermis (*F*), the average fibroblasts density in mm^3^ for all fields 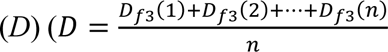 *where n is the number of observations*) was multiplied on mouse dermis volume (*V*).

### Histology, *in situ* hybridization, Immunohistology, and Second harmonic generation microscopy

Paraffin or OCT embedded skin tissues were stained with hematoxylin and eosin (H&E) or Masson’s trichrome by the standard procedures. For *in situ* hybridization, premade RNAscope™ probes for procollagen (Col1a1) and Yap/Taz, along with the necessary reagents, were purchased from Advanced Cell Diagnostics. The *in situ* hybridization procedure was carried out according to the manufacturer’s protocol. Immunohistology was performed as described previously (Quan et al. 2006), using antibodies against *Yap/Taz* (Cell Signaling Technology), type I procollagen (Santa Cruz Biotechnology). Briefly, paraffin embedded skin samples were sectioned (7 µm), permeabilized with 0.5% Triton X-100 in phosphate-buffered saline (PBS), blocked with rabbit serum (5% in PBS) and incubated for one hour at room temperature with primary antibodies, followed by incubation with secondary antibody for one hour at room temperature. After staining, the slides were examined using a digital imaging microscope (Zeiss). Specificity of staining was determined by substituting isotype-control immunoglobulin (mouse IgG2a) for the primary antibodies. No detectable staining was observed with isotype-controls. Second harmonic generation microscopy was performed using a Leica SP8 Confocal Microscope with 2-Photon, at the University of Michigan Microscopy and Image Analysis Laboratory.

### Mouse skin dermal fibroblasts isolation

The mouse skin was shaved, and any remaining hair was fully removed using a hair remover (Nail Lotion, Church & Dwight Co., Inc.). Then, the skin was washed with phosphate buffered saline (PBS) and sterilized with 70% ethanol twice. The skin tissue was placed in 60 mm tissue culture palate, then dissected and minced with surgical scissors (∼1mm) using sterile techniques under the hood. The minced tissue was incubated with 5ml of Liberase TL (Roche Diagnostics) solution (0.25mg/ml) at 37 C° temperature for one hours with gentle rotation. Tissue debris was removed by filtration through a 70 µm nylon Cell strainer (BD Biosciences), and the cells were collected by brief centrifugation. The cells were resuspended in Dulbecco’s modified Eagle’s minimal essential medium (DMEM; Life Technology Inc.) supplemented with 10% fetal bovine serum (FBS; Sigma Chemical Co.), penicillin (100 U/ml), streptomycin (100 μg/ml) in a humidified incubator with 5% CO_2_ at 37 C°.

### RNA isolation and quantitative real-time PCR

Total RNA was isolated from mouse skin or cultured cells using the RNeasy Mini Kit (Qiagen, Chatsworth, CA) following the manufacturer’s instructions. cDNA was synthesized from RNA using the TaqMan Reverse Transcription Kit (Applied Biosystems). Quantitative PCR was performed in duplicate using 2μl of cDNA per reaction with gene-specific primers (Real Time Primers, LLC), TaqMan Universal PCR Master Mix (Applied Biosystems), and a 7700 Sequence Detector System (Applied Biosystems). All PCR steps were carried out using a Biomek 2000 robotic workstation (Beckman Coulter, Inc.) to ensure accuracy and reproducibility. Target gene expression levels were normalized to the housekeeping gene 36B4.

### Western analysis

Whole cell proteins were extracted from dermal fibroblasts using WCE buffer (25 mM HEPES, 0.3 M NaCl, 1.5 mM MgCl₂, 0.2 mM EDTA, 1% Triton X-100, 20 mM β-glycerol phosphate). Protein expression was assessed by Western blot analysis. Briefly, proteins were separated on 10% SDS-PAGE gels, transferred to PVDF membranes, and blocked for 1 hour in PBST (0.1% Tween-20 in PBS) containing 5% non-fat milk. Membranes were incubated with primary antibodies for 1 hour at room temperature, washed three times with PBST, and then incubated with appropriate secondary antibodies for another hour at room temperature. After three additional washes with PBST, signal detection was performed using the ECF Western Blotting System (Amersham Pharmacia Biotech) according to the manufacturer’s instructions. Membranes were scanned using a STORM PhosphorImager (Molecular Dynamics). To assess loading consistency, membranes were stripped and reprobed with an anti-β-actin antibody (Sigma-Aldrich). Target protein intensities were quantified and normalized to β-actin levels.

### Transient transfection

Mouse dermal fibroblasts were transiently transfected with a YAP/TAZ mutant plasmid (Chan et al. 2011) by electroporation using dermal fibroblasts nucleofector kit (Amaxa Biosystems, Gaithersburg, MD). The YAP/TAZ mutant plasmid was generously provided by Dr. S.W. Chan (Institute of Molecular and Cell Biology, Agency for Science, Technology and Research, Singapore.

### RNA sequencing and spatial transcriptomics

RNA as isolated from the cells using RNeasy Micro kit (Qiagen, Chatsworth, CA), and the quality of the RNA was validated using 2100 Bioanalyzer. RNA sequence was performed by Novagen (Sacramento, CA) using the Illumina NovaSeq 6000 system (Illumina). Briefly, stranded mRNA libraries were prepared using the TruSeq RNA library prep kit (Illumina) and sequenced by Novagen. After library construction, the quality assessment was performed by Agilent 2100 Bioanalyzer and Q-PCR to accurately quantify the library effective concentration (> 2nM). Data were quality controlled and analyzed using standard pipeline for RNA-seq analysis, including adapter trimming, read mapping, and quantification of gene expression. Spatial transcriptomics was performed on paraffin-embedded skin tissue sections using the NanoString GeoMx® Digital Spatial Profiler (DSP), a platform designed for targeted transcriptomic analysis with spatial resolution. Regions of interest (ROIs) within the interstitial dermis, where dermal fibroblasts reside, were selected based on DAPI-stained images. Data were analyzed using GeomxTools v1.99.4. Differential gene expression between the two groups was analyzed using the linear mixed effect model (LMM). Pathway analysis was done using iPathwayGuide using all filtered gene targets as background. scRNA-seq dataset was generated with biopsies from paraffin-embedded mouse skin. Library preparation was carried out at the University of Michigan Advanced Genomics Core using the 10X Chromium system with v2 and v3 chemistry. Sequencing was performed on the Illumina NovaSeq 6000 platform, generating 150 bp paired-end reads. Quality control was performed by removing low-quality cells, which were defined as having fewer 200 genes, more than 5000 genes, or mitochondrial gene transcripts exceeding 20%. Ambient RNA was removed using SoupX (v1.6.2), and doublets were filtered out with scDblFinder (v1.12.0). Data normalization was performed with NormalizeData (“LogNormalize” method), followed by ScaleData and identification of the top 2000 variable genes using FindVariableFeatures. Dimensionality reduction was conducted via RunPCA on these variable genes, and UMAP was applied to the top 30 principal components (PCs). Batch effect correction was performed with Harmony (v1.0) using donor as the batch variable. Cells were clustered using FindNeighbors and FindClusters (resolution=0.5) with SNN modularity optimization. For visualization, RunUMAP was applied to the top 30 PCs. Cluster-specific marker genes were identified using FindAllMarkers in Seurat (v4.4.1), and cell types were assigned by comparing these markers to curated signature genes. Cells and clusters annotated as “Fibroblasts” were subsisted out for downstream analyses. Data processing, including quality control, read alignment to the hg38 reference genome, and gene quantification, was performed using the 10X Cell Ranger software suite. Subsequently, the samples were combined into a single expression matrix utilizing the Cell Ranger aggr pipeline.

### Sample preparation for proteomic analysis

Skin samples were processed as previously described (McCabe et al. 2025). Tissue samples snap-frozen in liquid nitrogen were lyophilized overnight using a FreeZone 4.5 L benchtop freeze dryer (Labconco #7750020) and milled to a fine powder prior to extraction. Three milligrams of material from each sample was combined with 100 mg of 1 mm glass beads (Next Advance #GB10) and homogenized in 200 μL/mg of decellularization buffer (50 mM Tris-HCl (pH 7.4), 0.25% CHAPS, 25 mM EDTA, 3 M NaCl) at power 8 for 3 min (Bullet Blender, Model BBX24, Next Advance, Inc.). Homogenate was then vortexed (power 8) for 20 min at 4°C, spun at 18,000 x g (4°C) for 15 min, and the supernatant was collected. Decellularization was repeated for a total of 3 washes, homogenizing and vortexing samples before each collection, and all washes were pooled to generate the cellular fraction. Pellets were then treated with freshly prepared hydroxylamine (HA) buffer (1 M NH_2_OH−HCl, 4.5 M Gnd−HCl, 0.2 M K_2_CO_3_, pH adjusted to 9.0 with NaOH) at 200 μL/mg of the starting tissue dry weight. Samples were homogenized at power 8 for 1 min and incubated at 45°C with shaking (1000 rpm) for 4 hours. Following incubation, the samples were spun for 15 min at 18,000 x g and the supernatant was removed before being stored at −80°C as the extracellular matrix (ECM) fraction until further proteolytic digestion with trypsin. All fractions were subsequently subjected to overnight enzymatic digestion with trypsin (1:100 enzyme:protein ratio) using a filter aided sample preparation (FASP) approach (Wisniewski et al. 2009) and desalted during Evotip loading (described below).

### LC-MS/MS analysis

Digested peptides (200 ng) were loaded onto individual Evotips following the manufacturer’s protocol and separated on an Evosep One chromatography system (Evosep, Odense, Denmark) using a Pepsep column, (150 µm inter diameter, 15 cm) packed with ReproSil C18 1.9 µm, 120Å resin. Samples were analyzed using the instrument default “30 samples per day” LC gradient. The system was coupled to the timsTOF Pro mass spectrometer (Bruker Daltonics, Bremen, Germany) via the nano-electrospray ion source (Captive Spray, Bruker Daltonics). The mass spectrometer was operated in PASEF mode. The ramp time was set to 100 ms and 10 PASEF MS/MS scans per topN acquisition cycle were acquired. MS and MS/MS spectra were recorded from m/z 100 to 1700. The ion mobility was scanned from 0.7 to 1.50 Vs/cm^2^. Precursors for data-dependent acquisition were isolated within ± 1 Th and fragmented with an ion mobility-dependent collision energy, which was linearly increased from 20 to 59 eV in positive mode. Low-abundance precursor ions with an intensity above a threshold of 500 counts but below a target value of 20000 counts were repeatedly scheduled and otherwise dynamically excluded for 0.4 min.

### Global proteomic data analysis

Data was searched using MSFragger v4.1 via FragPipe v22.0^2^. Precursor tolerance was set to ±12 ppm and fragment tolerance was set to ±25 ppm. Data was searched against UniProt restricted to *Mus musculus* reviewed sequences with added common contaminant sequences^3^ (17,340 total sequences). Enzyme cleavage was set to semi-specific trypsin for all samples. Fixed modifications were set as carbamidomethyl (C). Variable modifications were set as oxidation (M), oxidation (P) (hydroxyproline), deamidation (NQ), Gln->pyro-Glu (N-term Q), and acetyl (Peptide N-term). Label free quantification was performed using IonQuant v1.10.27 with match-between-runs enabled and default parameters. Cell and ECM fractions were searched separately and merged after database searching. Results were filtered to 1% FDR at the peptide and protein level.

### Statistical analysis

Statistical analysis was performed using GraphPad Prism (v.8) with unpaired two-sided Student’s t-tests, one-way analysis of variance (ANOVA) with Tukey’s method for multiple comparisons, or Kruskal-Wallis test with Dunn’s multiple comparisons test. Statistical significance was defined as P<0.05. All experiments were repeated a minimum of three times unless otherwise stated. Images were quantified by Image J software (Version 1.5p). All data are represented as Mean±SEM. Data Availability Statement

## ACKNOWLEDGMENTS

This work was supported by the National Institute of Health (RO1ES014697 to TQ, R01AG081805 and R01AG083378 to TQ, GJF).

## AUTHOR CONTRIBUTIONS

Conceptualization: TQ; Data Curation: AE, AJK, KCH, MM, JK, ZQ, ZZ, TH, CG, TQ; Formal Analysis: AE, AJK, KCH, MM, JK, ZQ, TQ; Investigation: AE, AJK, KCH, JK, ZQ, ZZ, TH, CG, TQ; Methodology: AE, TQ; Supervision: TQ, GJF; Visualization: AE, AJK, JK, ZQ, ZZ, TH, CG, TQ; Validation: AE, AJK, JK, ZQ, ZZ, TH, CG, TQ; Writing-Original Draft Preparation: TQ; Review and Editing: AE, AJK, KCH, MM, JK, ZQ, ZZ, TH, CG, GJF, JJV, TQ.

## DATA AVAILABILITY STATEMENT

The RNA-sequencing dataset related to this article can be found at https://www.ncbi.nlm.nih.gov/geo/, hosted at Home Geo National Center for Biotechnology Information (GSE305928). The data that support the findings of this study are available from the corresponding author upon reasonable request.

## Abbreviations

Yap: yes-associated protein
Taz: transcriptional coactivator with PDZ-binding motif
Col1: type I collagen
ECM: extracellular matrix

